# Multimodal meta-analysis of fecal metagenomes reveals microbial single nucleotide variants as superior biomarkers for early detection of colorectal cancer

**DOI:** 10.1101/2023.02.04.527090

**Authors:** Wenxing Gao, Xiang Gao, Lixin Zhu, Sheng Gao, Ruicong Sun, Zhongsheng Feng, Dingfeng Wu, Zhanju Liu, Ruixin Zhu, Na Jiao

## Abstract

Microbial signatures show remarkable potentials in predicting colorectal cancer (CRC). This study aimed to evaluate the diagnostic powers of multimodal microbial signatures, multi-kingdom species, genes, and single-nucleotide variants (SNVs) for detecting pre-cancerous adenomas. We performed cross-cohort analyses on whole metagenome sequencing data of 750 samples via xMarkerFinder to identify adenoma-associated microbial multimodal signatures. Our data revealed that fungal species outperformed species from other kingdoms with an area under the ROC curve (AUC) of 0.71 in distinguishing adenomas from controls, while classifier based on SNVs of bacterial species displayed the strongest diagnostic capability (AUC=0.89). SNV biomarkers also exhibited outstanding performances in three independent validation cohorts (AUCs = 0.83, 0.82, and 0.76, respectively) and were highly specific to adenoma. In further support of the above results, functional analyses revealed more frequent inter-kingdom associations between bacteria and fungi, and abnormalities in quorum sensing, purine and butanoate metabolism in adenoma, which were further validated in a newly-recruited cohort via qRT-PCR. Therefore, these data extend our understanding of adenoma-associated multimodal alterations in the gut microbiome and provide a rationale of microbial SNVs for the early detection of CRC.

## Introduction

Colorectal cancer (CRC), currently the second most frequently diagnosed cancer, accounts for approximately 10% of all new cancer cases globally^1^. Recent data reveal a rising incidence of CRC in individuals aged under 50 years^2^, indicating a heavier burden in the coming years for the healthcare system worldwide. Of note, pre-cancerous adenoma is a major precursor form of CRC about 10-15 years preceding cancer initiation, the early detection and removal of which could significantly alleviate the incidence and mortality of CRC^1,3^.

A wide variety of strategies are available for CRC diagnosis, including the combination of colonoscopy and histopathology as the gold standard, and non-invasive test kits such as fecal occult blood test and fecal immunochemical test^4^. Progresses have also been made in blood or stool-based biomarkers, such as circulating tumor cells (CTC), circulating tumor DNA (ctDNA) and exosomes, broadly applied as non-invasive tools for tumor diagnosis, and for predictions of tumor recurrence and metastasis^5,6^. However, these approaches are dissatisfactory with high false-positive rates (up to 76.4%) and poor sensitivity (down to 38%) for adenoma^7,8^. Thus, there is an urgent need to explore and identify novel biomarkers specifically targeting the pre-cancerous adenoma stage for the purpose of early detection of CRC.

As a frequently used proxy of intestinal microbiome, fecal sample has demonstrated some potentials for detecting CRC in diagnostic models based on bacterial abundances^9–11^. For early screening of CRC, bacterial species achieved an area under the ROC curve (AUC) of 0.80 for diagnosing adenomas with multiple cohorts^12^. Besides bacteria, the previously neglected non-bacteria microorganisms, such as fungi, archaea and viruses, are gaining attentions as novel candidate disease biomarkers. Alterations in non-bacterial enteric microbiome and of intra- and inter-kingdom microbial interactions have also been revealed in non-alcoholic fatty liver disease (NAFLD)^13^, inflammatory bowel disease (IBD)^14^ and CRC^15–18^. Along this direction, we recently achieved improved specificity and accuracy for early-stage CRC screening with microbial multi-kingdom species compared to single-kingdom species^19^. On the other hand, it has been reported that CRC associates with microbial genes more robustly than with microbial species,^20^ reflecting the importance of the functional omics in health and disease^21^. Yet another type of microbial features, single nucleotide variants (SNVs), has emerged as effective diagnostic biomarkers for CRC and other diseases^22,23^. As such, one immediate question is, what are the predictive capabilities of the microbial multi-kingdom species, genes, and SNVs for pre-cancerous adenomas, and whether the combination of different types of microbial features could outperform the individual type of microbial features?

To address these questions, we performed comprehensive analyses on whole metagenome sequencing (WMS) data from seven cohorts (750 samples) to systematically explore the capability of multimodal biomarkers for detecting adenoma, aiming to facilitate the early detection of CRC. We observed that the SNV-based diagnostic model achieved superior accuracy (AUC=0.89), drastically outperforming species- and gene-based models for adenoma diagnosis. Furthermore, functional dysbiosis related to microbial quorum sensing, purine and butanoate metabolism was observed in the microbiome of adenoma patients.

## Results

### Characteristics of multiple cohorts and consistent processing of metagenome data

In this study, we included fecal WMS data from six published studies and one in-house cohort with samples collected in China to establish universal microbial multimodal biomarkers for adenoma (Fig. 1a, Data S1). There were in total 750 samples including 183 colorectal adenoma patients and 439 healthy controls in the discovery dataset, as well as 63 adenoma patients and 64 healthy controls in the external validation dataset. In addition, 59 samples (29 adenoma patients and 30 healthy controls) were newly collected in China to conduct qRT-PCR validations.

**Fig. 1.**
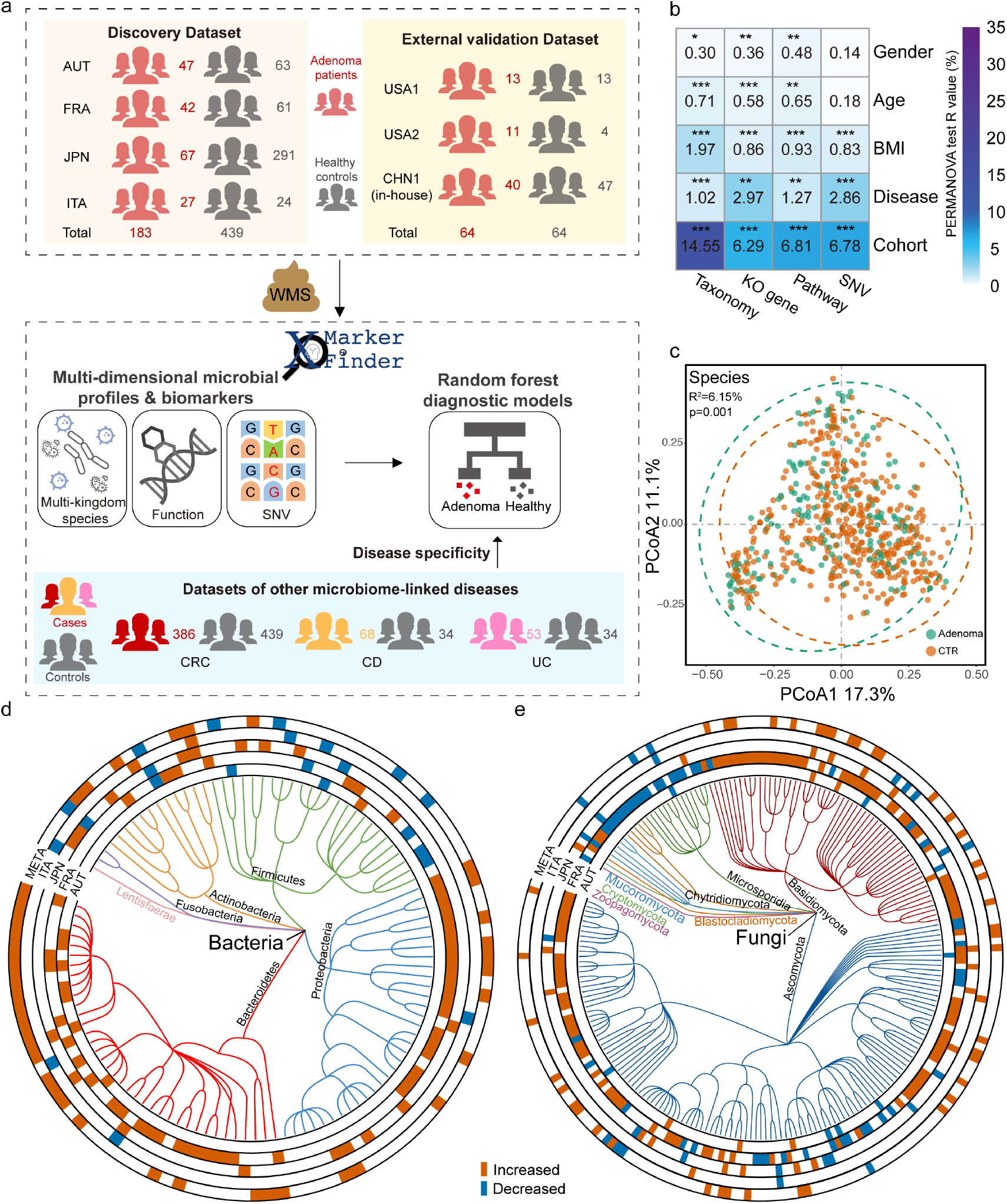
Experimental design and the integrated analysis of adenoma-associated microbiome. **a**, Experimental design. A total of 622 samples comprising 183 colorectal adenoma patients and 439 healthy controls were included as the discovery dataset. WMS data were then processed consistently to establish cross-cohort multimodal biomarkers and diagnostic models via xMarkerFinder. Further, the external validation dataset (128 samples) and datasets of other microbiome-linked diseases were used to independently assess the robustness and disease specificity of established biomarkers and diagnostic models. **b**, The PERMANOVA test identified “cohort” as the major confounder and demographic indices (gender, age, and BMI) as minor confounders. Asterisks: statistical significance (*, *P*< 0.05; **, *P*< 0.01;***, *P*< 0.001). **c**, Principal coordinate analysis (PCoA) of microbial taxonomic classifications showing that gut microbiota differed between adenoma patients and healthy controls (*P*= 0.001). P values of beta diversity based on Bray-Curtis distance were calculated with PERMANOVA test. Each point in the PCoA plots represents a sample and the colors of points represent different groups. **d**, **e**, Phylogenetic trees showing the differential bacterial (**d**, 110 in total) and fungal (**e**, 216 in total) species. The outer circles are marked for significant differential species (*P* < 0.05) in each cohort identified via Maaslin2 and in the meta-analysis identified via MMUPHin (META ring). Orange and blue indicate increased and decreased abundance of the species, respectively.

We performed an integrated analysis of metagenome-derived microbial taxonomic, functional, and SNV profiles. To avoid technical bias across cohorts, all raw sequencing data were reprocessed consistently for microbial multimodal profiling. Considering the heterogeneity across different cohorts, the effects of major metadata variables on profiles of each modality were estimated by Permutational multivariate analysis of variance (PERMANOVA) test, which revealed the predominant impact of “cohort” explaining the largest proportions of variances in all data layers (Fig. 1b). Following “cohort”, disease status, body mass index (BMI), age, and gender exhibited lower but significant impact on the microbial profiles. Therefore, “cohort” was treated as the major confounder while gender, age, and BMI used as covariates in the identification of multimodal differential signatures by Meta-analysis Methods with a Uniform Pipeline for Heterogeneity in Microbiome Studies (MMUPHin) wrapped in xMarkerFinder.

### Adenoma-associated microbial multi-kingdom species

With all samples from the discovery cohorts, beta diversity based on Bray-Curtis dissimilarity among the taxonomic profiles showed that gut microbiota highly differd between adenoma patients and healthy controls (R^2^= 6.15%, *P*= 0.001, Fig. 1c). On the other hand, a trend of increased alpha diversity was observed in adenoma patients than that of controls, albeit not statistically significant (Fig. S1a, b).

We then explored adenoma-associated microbial taxonomic signatures. As expected, different sets of bacterial species were identified as differential species in distinct cohorts (Fig. 1d, e, Fig. S1c, d), necessitating the integrated analysis on demographically distinct populations. With the combined discovery dataset, we identified 46 differential bacterial species between adenoma and control (Data S2, Fig. 1d). Consistent with our previous work^12^, six bacterial species with decreased abundances in adenoma were observed, including *Bifidobacterium longum, Ruminococcus bicirculans, Longibaculum sp. KGMB06250, Eggerthella lenta, Blautia sp. YL58*, and *Enterococcus faecium*. Meanwhile, the abundances of 40 bacterial species were increased in adenoma compared with control, including *Alistipes shahii, Paraprevotella xylaniphila, Bacteroides helcogenes*, and particularly, two pathogenic bacteria, *Bacteroides caccae* and *Prevotella intermedia*.

Next, we examined the alterations in non-bacterial microorganisms including fungi, archaea and viruses. The abundances of 42 out of 50 differential fungal species were increased in adenoma compared to control, including *Sistotremastrum suecicum, Postia placenta, Kwoniella bestiolae*, and *Fusarium pseudograminearum*, while the abundances of *Rhizophagus irregularis, Aspergillus niger, Aspergillus ochraceoroseus, Leucoagaricus sp. SymC.cos, Aspergillus japonicus, Hyphopichia burtonii, Nematocida parisii*, and *Enterospora canceri* were decreased (Data S2, Fig. 1e). For archaea, among 11 significantly differential archaeal species, the abundances of *Thermococcus eurythermalis* and *Methanothrix soehngenii* were increased, while other nine archaeal species were decreased in adenoma patients (Data S2, Fig. S1c). For viruses, six differential signatures were identified. *Enterobacteria phage P4, Salmonella phage epsilon34*, and giant viruses including *Pandoravirus inopinatum* and *Orpheovirus IHUMI-LCC2* were observed with greater abundances in adenoma compared to control, while *Mycobacterium virus Giles* and *Escherichia virus RB49* were with lower abundances (Data S2, Fig. S1d). These microbial multi-kingdom signatures presented consistent changes in adenoma across cohorts.

### Adenoma-associated microbial functional alterations

Microbial functional alterations were examined at KEGG orthology (KO) gene and pathway levels, respectively. Significant differences in the beta diversity analysis of the entire set of microbial KO genes between adenoma and control (R^2^=2.97%, p=0.014, Fig. 2a) indicated altered microbial functions in adenoma. On the other hand, increased alpha diversity of the microbial genes was observed in adenoma patients compared to controls (Fig. S2 a, b).

**Fig. 2.**
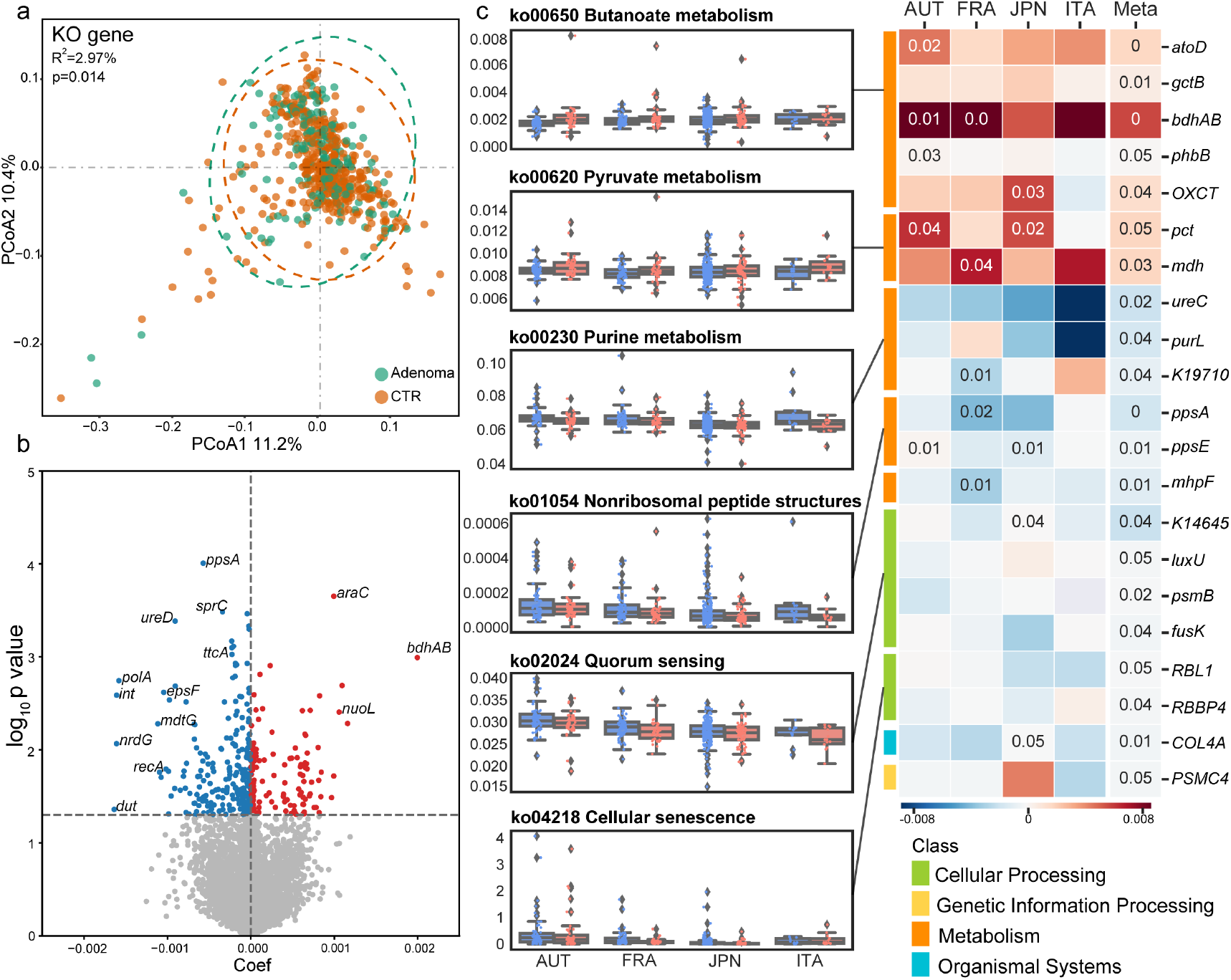
Adenoma-associated microbial functional alterations. **a**, PCoA of microbial functional KO genes showing that gut microbiota differed between adenoma patients and healthy controls (p=0.014). *P* values of beta diversity based on Bray-Curtis distance were calculated with PERMANOVA. Each point in the PCoA plots represents a sample and the colors of points represent different groups. **b**, Volcano plot showing the differential KO genes in all samples identified via MMUPHin. Each point represents a KO gene. Coefficient > 0, *P* < 0.05 (in red): genes significantly more abundant in adenoma compared with control; Coefficient < 0, *P* < 0.05 (in blue): genes significantly less abundant in adenoma compared with control; *P*> 0.05 (in grey): non-differential genes. **c**, The box plots (left) show the relative abundances of pathways in adenoma (red) and control (blue). Heatmap (right) shows the abundances of relevant differential KO genes in each of the four cohorts. Coefficient values were calculated via MaAsLin2 in each cohort and MMUPHin in meta-analysis with red for down-expression and blue for over-expression in adenoma patients compared with healthy controls. Only *P*-values < 0.05 are shown in the cells.

Similarly, the integrated analysis identified 386 differential KO genes, including 150 KO genes with increased abundances and 236 KO genes with decreased abundances in adenoma patients compared with healthy controls (Fig. 2b, Data S3). At pathway level, 15 differential pathways were identified (Fig. 2c, Data S4). Among these, five pathways enriched in adenoma were related to metabolism, such as butanoate metabolism, pyruvate metabolism, and styrene degradation, and organismal systems, including proximal tubule bicarbonate reclamation and transduction olfactory. Consistently, KO genes related to above pathways were more abundant in adenoma, such as propionate CoA-transferase (*pct*), and malate dehydrogenase (*mdh*) in pyruvate metabolism, as well as acetate CoA/acetoacetate CoA-transferase alpha subunit (*atoD*), glutaconate CoA-transferase subunit B (*gctB*), butanol dehydrogenase (*bdhAB*), acetoacetyl-CoA reductas*e (phbB*) and 3-oxoacid CoA-transferase (*OXCT*) in butanoate metabolism (Fig. 2c). On the other hand, 10 pathways and relevant KO genes were negatively associated with adenoma. These pathways and KO genes were mainly involved in metabolism and cellular processes (Fig. 2c). Specifically, the pathway of quorum sensing, a microbial cell-to-cell communication mechanism, was less represented in adenoma. Consistently, relevant KO genes, mainly genes of two-component systems, such as phosphorelay protein (*luxU*), and sensor histidine kinase *fusK*) exhibited lower abundances in adenoma group. Similarly, purine metabolism pathway and corresponding genes, phosphoribosylformylglycinamidine synthase (*purL*), and urease subunit alpha (*ureC*)were less abundant in adenoma patients (Fig. 2c). Collectively, these analyses revealed global alterations in microbial genes and pathways in adenoma across multiple cohorts.

### Adenoma-associated microbial SNV signatures

The microbial genetic variation, SNV, represents potential alterations in sub-genome information of microbial functionality. We next examined the SNV signatures of adenoma against 28 commonly detected microbial strains in the discovery dataset. The numbers of SNVs varied greatly, ranging from 56 in *Streptococcus salivarius* to 154089 in *Faecalibacterium prausnitzii* (61481) (Fig. 3a, Data S5). Among these, four strains belonged to above identified differential species, including *A.shahii* (24945 SNVs) and *B.caccae* (35193 SNVs) with greater relative abundances in adenomas compared with controls and *R.bicirculans* (96667 SNVs) and *B.longum* (43920 SNVs) with lower relative abundances (Data S5).

**Fig.3.**
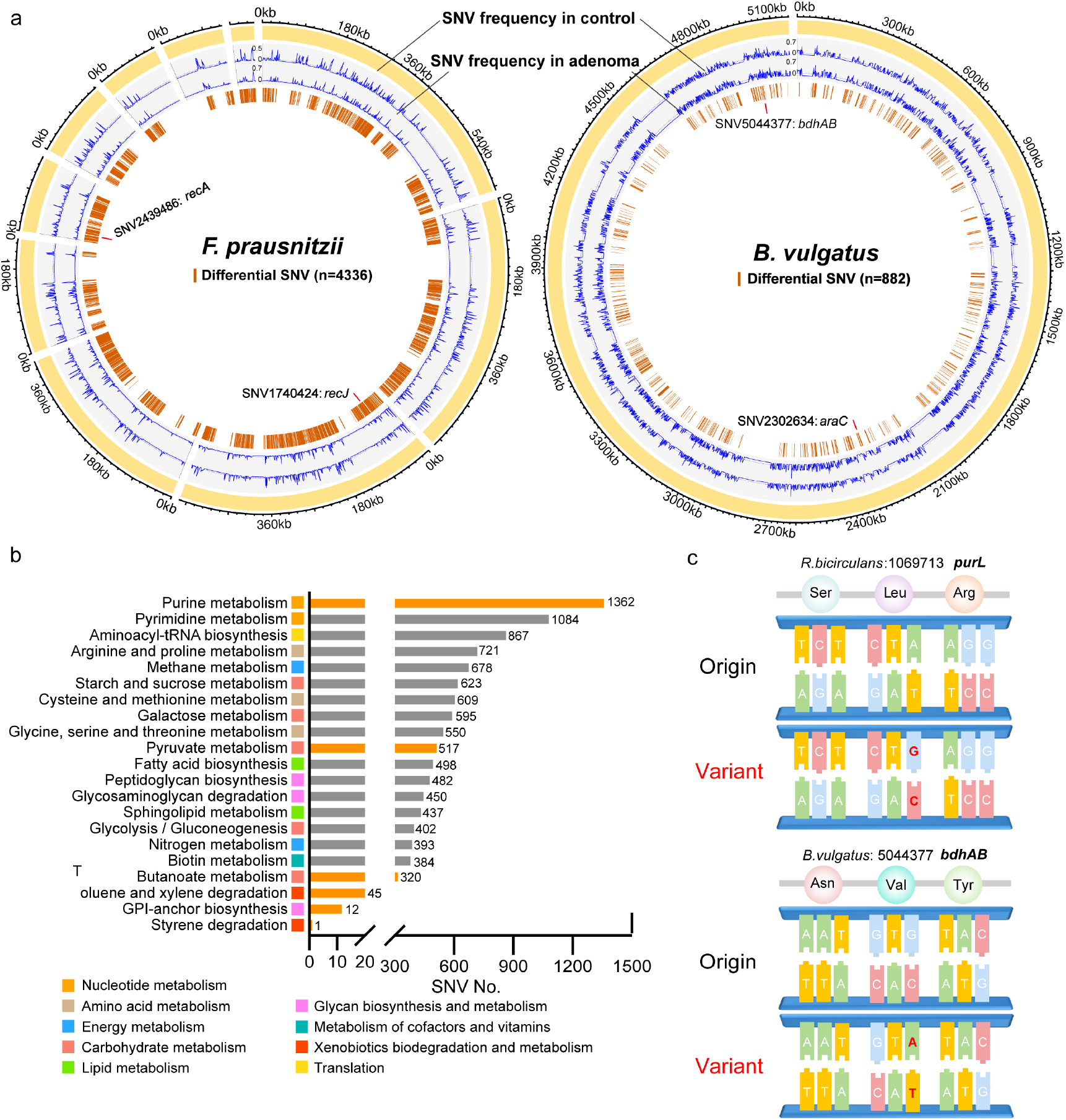
Microbial SNV signatures in adenoma. **a**, Genomic locations of SNVs in the strains of *F. prausnitzii* (61481) and *B.vulgatus* (57955). Yellow outer rings represent the genomes of the annotated strains. SNV frequencies in adenoma and control were indicated by blue lines in the 2^nd^ and 3^rd^ rings, respectively. The 4^th^ rings indicate locations of identified differential SNVs (brown lines). **b**, Bar plots showing the number of differential SNVs that belongs to each functional pathway with orange indicating previously established differential pathways. **c**, Mapping of two differential SNVs located in differential genes. Mutated nucleotides (red) and corresponding amino acids were shown.

Meta-analytic differential testing identified SNVs with consistently differential frequencies between adenoma and control samples across four cohorts. The numbers of differential SNVs annotated from one single strain ranged from 17 to 4658, with an exception of *S. salivarius*, of which no differential SNVs were identified (Data S5). Among them, *B.caccae* and *Parabacteroides distasonis* from the Bacteroidetes phylum and *F. prausnitzii*, *R.bicirculans* and *Roseburia intestinalis* from the Firmicutes phylum provided most of the differential SNVs, with an average number of 4034 (Data S5). Most of the differential SNVs were located in coding sequence (CDS) with only 10% in intergenic region (IGR). Meanwhile, differential SNVs were mainly located in metabolism-related genes (Fig. 3b), especially genes of purine metabolism (1362 SNVs) and pyrimidine metabolism (1084 SNVs) of nucleotide metabolism pathways, as well as amino acid metabolism pathways, including arginine and proline metabolism (721 SNVs), cysteine and methionine metabolism (609 SNVs) and glycine, serine and threonine metabolism (550 SNVs). Notably, considerable differential SNVs were related to the aforementioned differential functional pathways, such as purine metabolism, pyruvate metabolism (517 SNVs), butanoate metabolism (320 SNVs) and styrene degradation (one SNV). Additionally, several differential SNVs were located in previously identified differential genes, such as SNV (1069713) of *R. bicirculansin* in *purL* and SNV (5044377) of *Bacteroides vulgatus* in *bdhAB* (Fig. 3c). These findings further underlined the importance of microbial genetic variations in the pathology of adenoma and the potential diagnostic capabilities of SNVs.

### The diagnostic models for adenoma based on microbial multimodal biomarkers

The performance of diagnostic models highly depends on features. Thus, to determine optimal features for model construction, we employed Triple-E, a comprehensive three-step feature selection procedure in xMarkerFinder, which mainly comprises feature effectiveness evaluation, collinear feature exclusion and recursive feature elimination (Fig. 4a, detail in Methods). Furthermore, random forest (RF) models were optimized via adjustment of hyperparameters.

**Fig. 4.**
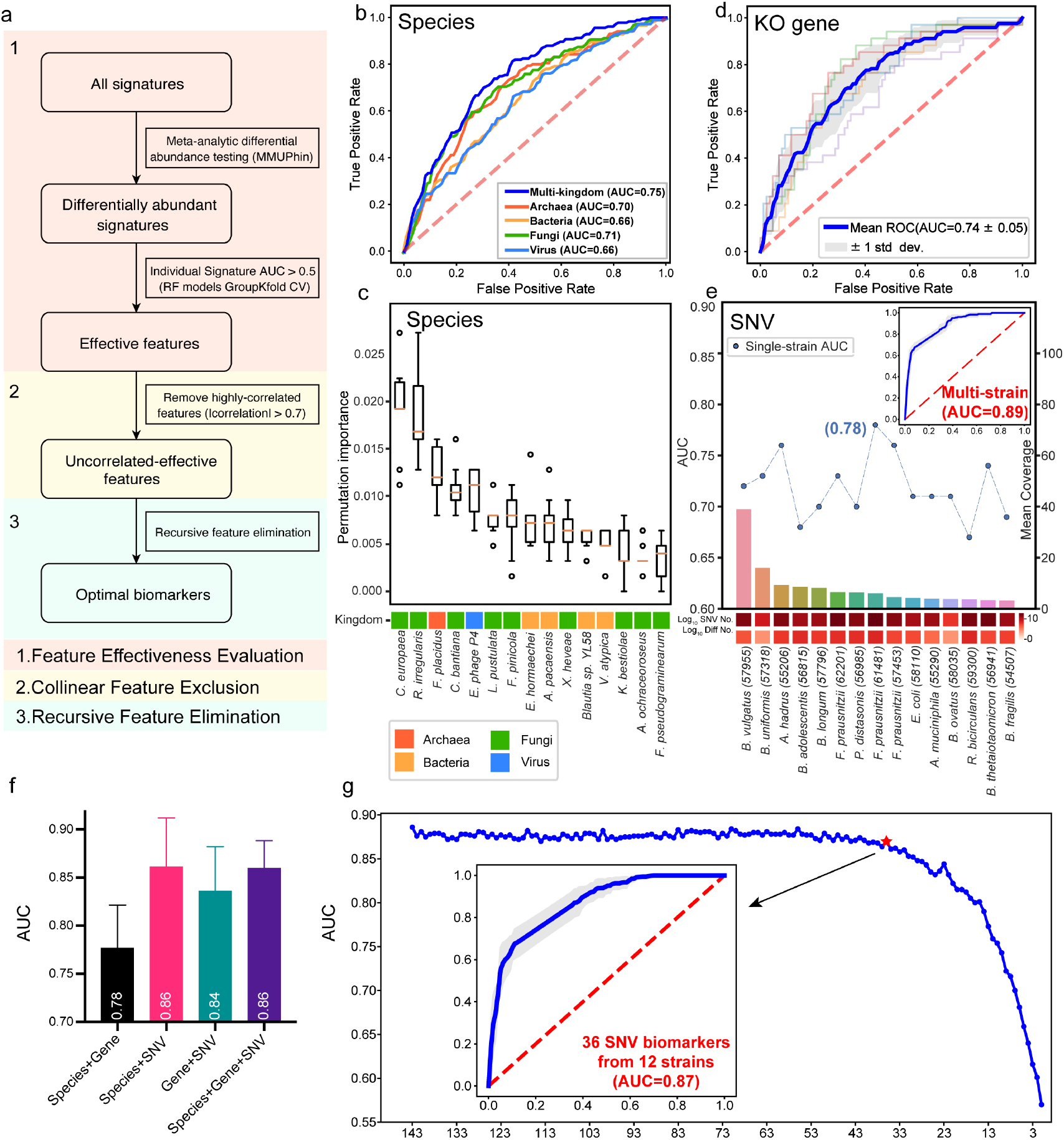
Diagnostic models based on microbial multimodal biomarkers. **a**, The workflow of “Triple-E” feature selection procedure in xMarkerFinder. (1) Feature Effectiveness Evaluation. The differentially abundant signatures were identified using MMUPHin out of all multimodal signatures. Each single differential signature was then used to build a RF model with GroupKfold cross-validation and signatures with AUC values above 0.5 were defined as “effective features”. (2) Collinear Feature Exclusion. For all effective features, only those with absolute values of Spearman’s rank correlation coefficients less than 0.7 were reserved as “uncorrelated-effective features”. (3) Recursive Feature Elimination. The recursive feature elimination method was utilized to determine the “optimal biomarkers” as the best panel of features used for model construction. **b**, Receiver operating characteristic (ROC) curve of the optimized models constructed with species-level biomarkers. Mean AUC and standard deviation of stratified 5-fold cross-validation were shown. **c**, Permutation feature importances of the optimized model constructed with multi-kingdom species-level features. Color represents different kingdoms. **d**, ROC curve of the optimized models constructed with KO gene-level biomarkers. Mean AUC and standard deviation of stratified 5-fold cross-validation were shown. **e**, The upper plot showing the performances of single-strain SNV models and the multi-strain SNV model. Bar plot shows the mean coverage of each strain across all samples in the discovery dataset. The log-transformed numbers of annotated SNVs and differential SNVs in each strain are color-coded and indicated below the bar plot. **f**, Box plot showing the cross-validation AUC values of models constructed with the combination of multimodal biomarkers. Mean AUCs and standard deviations were shown. **g**, The process of selecting the minimal panel of SNV biomarkers with the inner plot showing the ROC curve of the minimal panel of SNV biomarkers.

### Models based on multi-kingdom species outperform models with single-kingdom species

Diagnostic models based on bacterial abundances are widely used approaches for disease screening^9,12,24^. Considering the alterations in all four microbial kingdoms, we assessed the prediction capabilities of microbial features in four kingdoms, respectively. Unexpectedly, the diagnostic model constructed with eight fungal species displayed the strongest ability to distinguish adenoma from control (AUC=0.71), superior to models based on archaeal species (AUC=0.70, four biomarkers), bacterial species (AUC=0.66, five biomarkers), and viral species (AUC=0.66, four biomarkers, Fig. 4b). The AUC values of models based on two-kingdom and three-kingdom species features were slightly improved compared to those with single-kingdom features, ranging from 0.66 to 0.74 (Fig. S3), suggesting additive predictive value of the combination of species biomarkers from different kingdoms. Notably, the diagnostic model constructed with a core set of optimal species biomarkers from all four kingdoms achieved the highest AUC of 0.75 for distinguishing adenoma patients from controls (Fig. 4b). The observation that the best-performing multi-kingdom diagnostic model encompassed nine fungal species biomarkers out of a total of 15 species, including *Cyphellophora europaea* and *R. irregularis* acting as the most contributing biomarkers, further corroborated the prominent potential of the fungal kingdom in early detection of CRC (Fig. 4c).

### Microbial functional biomarkers for adenoma prediction

Functional classifiers were constructed with the differential genes and pathways, respectively. The optimal gene-based model comprising 31 KO genes achieved an AUC of 0.74, higher than those of the models constructed with single-kingdom species biomarkers while slightly lower than that of the best-performing multi-kingdom species model (Fig. 4d). Genes involved in metabolic pathways, such as prepilin peptidase (*pilD*), ethanolamine utilization protein (*eutN*), and *bdhAB*, and genes involved in genetic information process, such as putative DNA relaxase (*nicK*) and urease accessory protein (*ureE*), contributed most to the diagnostic capability of the model (Fig. S4). In addition, the diagnostic model based on seven optimal pathways achieved a relatively low AUC of 0.68 (Fig. S5), which may be rationalized by the fact that functional pathways provide aggregated gene information, thus neutralizing individual signatures’ variations.

### Microbial SNVs are better diagnostic biomarkers for adenoma

Given that microbial genetic variations have profound impact on its function and that the integrated analysis identified considerable consistent differential SNVs across cohorts, we next evaluated the diagnostic potential of microbial SNVs. To achieve high robustness and generalizability of SNV-based models, only strains with sufficient mean coverage across samples (> 3X) were preserved (Data. S5). Firstly, we identified optimal SNV biomarkers from each strain that achieved AUC values ranging from 0.67 to 0.78 (Fig. 4e, Data S5). Of note, three out of 15 single-strain SNV models outperformed both multi-kingdom species and gene models, including *F. prausnitzii*(61481) (AUC=0.78), *F. prausnitzii* (57453) (AUC=0.76), and *A. hadrus* (55206) (AUC=0.76, Fig 4e). The observation that the taxonomic abundances of *F. prausnitzii* and *A. hadrus* did not differ between adenoma and control further testified for the prevailing diagnostic capabilities and sensitivity of microbial SNVs in detecting adenoma. Furthermore, to evaluate the predictive ability of the combination of multistrain SNVs, optimal SNV biomarkers from each strain were pooled together and a multi-strain SNV model was established achieving a highest AUC value of 0.89 with a total of 143 SNV biomarkers (Fig 4e).

To better understand the multi-stain SNV diagnostic model, we examined the biomarkers’ distribution and found that 61 of the 143 SNV biomarkers were located in the genome of *F. prausnitzii* (8 in strain *62201*, 10 in strain *57453*, and 43 in strain *61481*), one of the most abundant species in human intestines (Data S6)^25^. 131 out of the 143 SNVs were located in CDS with nearly half (62 SNVs) being synonymous mutations that commonly do not alter the structure or function of the encoded proteins.

To test the hypothesis that the combination of multimodal biomarkers might provide additive predictive capabilities, we then evaluated the predictability of models based on the combination of optimal biomarkers from species, genes and SNVs. Improved performance (AUC=0.78) was observed with the combination of species- and genelevel biomarkers (Fig. 4f). However, we were surprised by the decreased diagnostic performances when adding species or gene biomarkers to the SNV model (Fig. 4f). Thus, the SNV model was the best-performing diagnostic model for detecting adenoma.

Considering the efficiency and cost effectiveness in clinical practice, we further identified a minimal set of SNV biomarkers via recursive feature elimination and the classifier constructed with a core set of 36 SNV biomarkers reached an AUC of 0.87 (Fig. 4g, 5a), outperforming the combination of multimodal biomarkers (AUC = 0.86). The core set of 36 SNV biomarkers were therefore considered the best panel of adenoma biomarkers for potential clinical application.

**Fig. 5.**
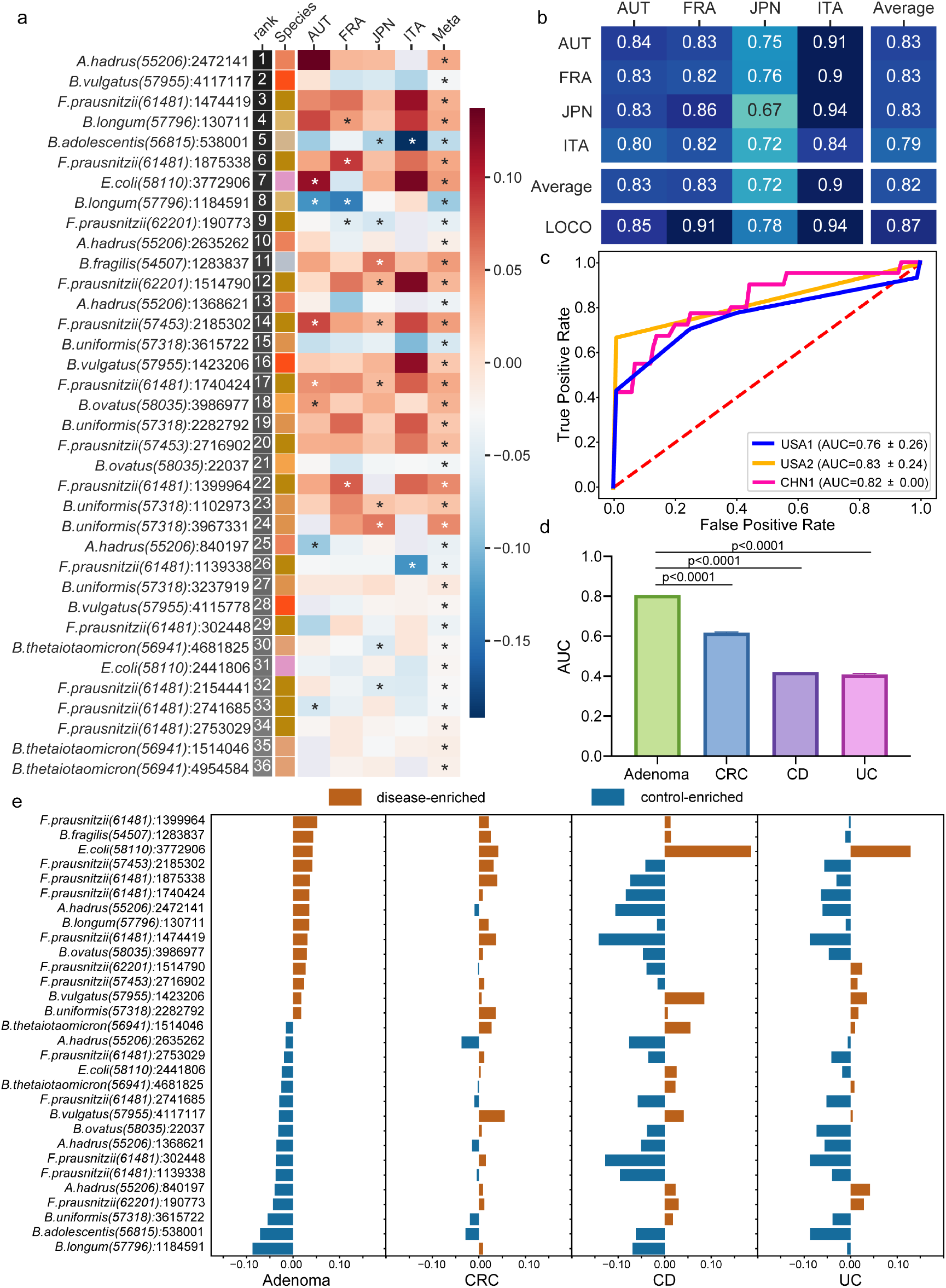
Characterization and validation of the best panel of SNV biomarkers. **a**, SNV biomarkers’ distribution in each cohort and in meta-analysis. Heatmap of the coefficient values calculated via MaAsLin2 in each cohort or via MMUPHin in meta-analysis, with red for higher frequencies and blue for lower frequencies in adenoma patients compared with healthy controls. Asterisks indicate statistical significances (*P*< 0.05). Biomarkers are ordered according to their permutation importances in the diagnostic model (Rank column). The Species column indicates different species where the SNVs were located (color coded). **b**, Internal validation AUC matrix. Values on the diagonal refer to the average AUC values of 5-fold cross-validation within each cohort. Off-diagonal values refer to the AUC values obtained by training the classifier on the cohort of the corresponding row and applying it to the cohort of the corresponding column. The LOCO row refers to the performances obtained by training the model using all but the cohort of the corresponding column and applying it to the cohort of the corresponding column. **c**, The performances of the optimal SNV biomarker panel in three external validation datasets. **d**, The boxplots showing the comparison of the performances of SNV biomarkers in RF models for different microbiome-linked diseases: adenoma, CRC, CD, and UC. *P* values were from two-sided Wilcoxon rank-sum tests. **e**, Patterns of the correlations of SNV biomarkers with disease status: distinct patterns observed for adenoma and other microbiome-linked diseases. Coefficient values calculated by MaAsLin2 and MMUPHin are plotted by color gradients with orange for disease-enriched SNVs and blue for control-enriched SNVs.

### Microbial SNV biomarkers are robust and universal across cohorts

To ascertain the reproducibility of the best panel of SNV biomarkers, cohort-to-cohort and leave-one-cohort-out (LOCO) validations were performed. The AUC values of the models constructed with the best panel of SNV biomarkers ranged from 0.67 to 0.94 with an average of 0.82 for cohort-to-cohort validation and ranged from 0.78 to 0.94 with an average of 0.87 in LOCO validation (Fig. 5b). To further test the robustness of the established best SNV panel, we evaluated their diagnostic capabilities with three external cohorts from different geographic regions and achieved AUC values of 0.83, 0.82 and 0.76, respectively (Fig. 5c). These substantial validations confirmed the robustness and generalizability of the identified best panel of microbial SNV biomarkers for adenoma diagnosis.

### Specificity of microbial SNV biomarkers for adenoma prediction

To evaluate the disease specificity of the best panel of SNV biomarkers, the diagnostic capabilities of this SNV model were evaluated for prediction of other microbiome-linked diseases, including CRC, Crohn’s disease (CD) and ulcerative colitis (UC). The predictive AUC values of the SNV model for these three diseases were apparently lower than that of adenoma (Fig. 5d), indicating a high specificity of SNV biomarkers for adenoma.

Additionally, we evaluated the correlations between each SNV biomarker and the disease status in adenoma and three non-adenoma diseases, and observed distinct patterns for adenoma and non-adenoma diseases (Fig. 5e). Importantly, in CRC compared to adenoma, these SNVs were less significantly correlated with disease status, reflecting the capabilities of these SNVs in distinguishing adenoma from CRC, or, the CRC early detection at pre-cancerous stage (Fig. 5e). CD and UC, the two main subtypes of IBD, shared similar SNV correlation patterns which differed from those of adenoma (Fig. 5e). Altogether, these analyses together demonstrated the disease specificity of the microbial SNV biomarkers for adenoma diagnosis.

### Cross-modality associations among the microbial signatures are indicative of pathogenic mechanisms in adenoma

To explore the potential mechanisms for the microbial multimodal signatures to participate in adenoma pathogenesis, we constructed the microbial co-abundance networks in adenoma patients and healthy controls and observed distinct community structures. The microbial network of adenoma patients (76 species, 259 associations) was far more complex than that of healthy controls (52 species, 185 associations, Fig. 6a). Notably, adenoma network contained more frequent intra- and inter-kingdom connections, such as interactions among intra-fungal species, *C. europaea-Thermothelomyces thermophilus, Cladophialophora bantiana-Ustilago maydis*, and *Aureobasidium melanogenum-K.bestiolae*, and fungal-baterial interactions among *H. burtonii-A. shahii, Aspergillus versicolor-Prevotella enoeca* and *Jaminaea rosea-Sorangium cellulosum*, indicating fungal involvement in adenoma pathogenesis. Moreover, eight species biomarkers from the fungal and bacterial kingdoms, including *C. europaea, C. bantiana, K. bestiolae, Lasallia pustulata, Fomitopsis pinicola, Enterobacter hormaechei, Actinomyces pacaensis* and *Blautia sp. YL58*, presented more associations than other species, suggesting a core set of microbiota contributing to adenoma pathogenesis.

**Fig.6.**
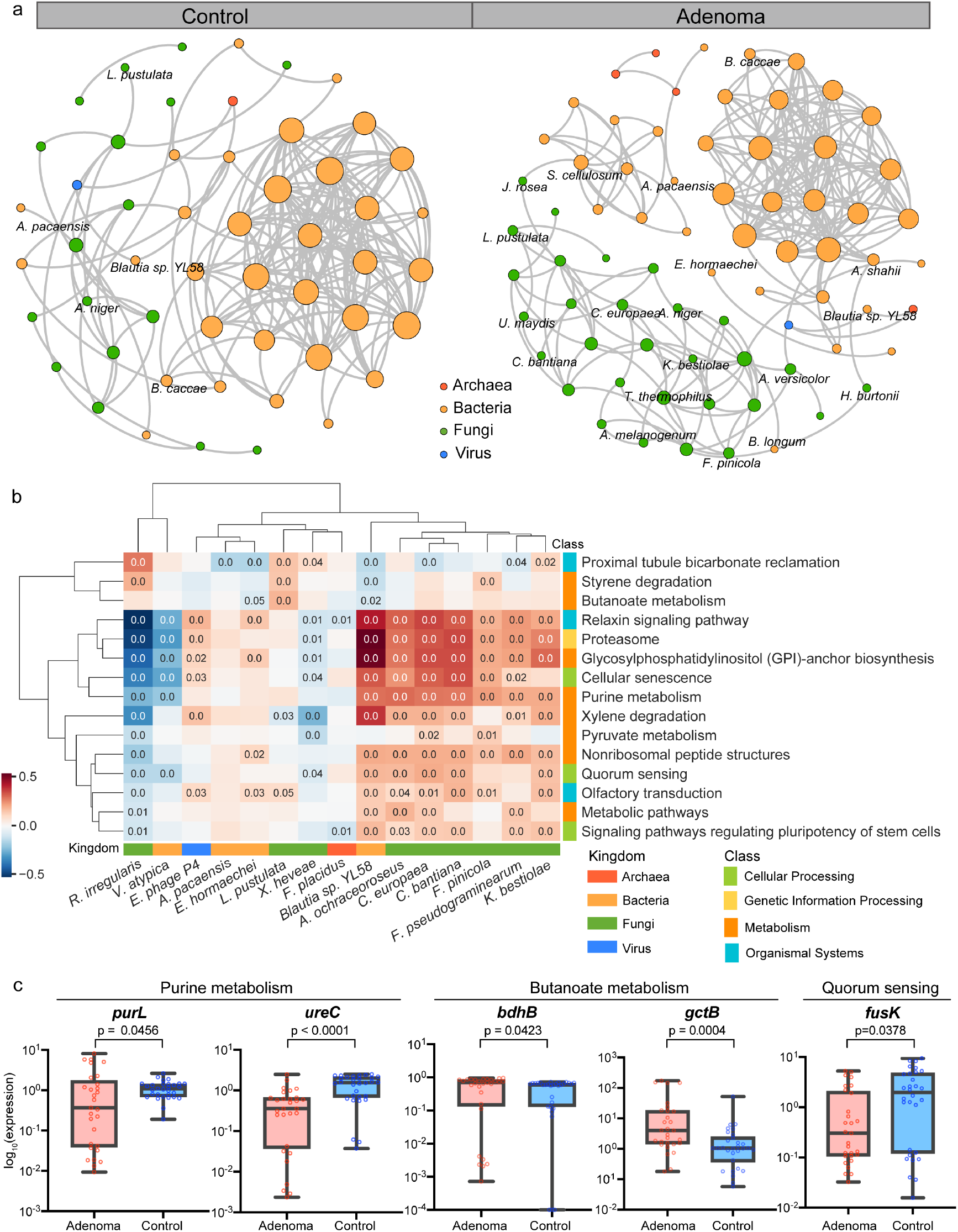
Cross-modality associations among the microbial signatures. **a**, Multi-kingdom co-abundance networks in adenoma and control. Colors of nodes indicate different kindoms: archaea (red), bacteria (yellow), fungi (green) and viruses (blue). Only correlations with *P*-values < 0.05 and absolute correlations coefficients > 0.5 are included in the networks. **b**, HAllA results showing associations between differential pathways and multi-kingdom species biomarkers. Significant associations between pathways and species were plotted (*P*< 0.05). Correlation coefficients are coded in color: red and blue indicate positive and negative correlations, respectively. Only p values less than 0.05 are shown in the cells. **c**, qPCR results showing the expression level of key genes in purine metabolism, butanoate metabolism, and quorum sensing.
*P* values were from two-sided Wilcoxon rank-sum tests.

Besides, considerable associations were observed between differential functional pathways and species, as well as the optimal multi-kingdom species biomarkers (Fig. S6, Fig. 6b). Specifically, the species-level biomarkers *A. ochraceoroseus*, *C. bantiana*, *C. europaea* and *K. bestiolae* of fungal kingdom and *Blautia sp. YL58* of bacterial kingdom were positively associated with pathways of purine metabolism and quorum sensing. Counter-intuitively, the bacterial biomarkers *E. hormaechei* and *Blautia sp. YL58*, two of the butyrate-producing species, were negatively associated with butanoate metabolism. Interestingly, one of the most important fungal biomarkers, *R. irregularis*,and bacterial biomarker *V. atypica* displayed rather distinct patterns from other species-level biomarkers and were negatively correlated with most differential functional pathways.

As described above, the abundances of key genes in these pathways displayed consistent alterations, such as decreased abundances of two-component system *luxU, fusK* in quorum sensing pathway, *ureC*, *purL* in purine metabolism and increased abundances of *atoD, gctB, bdhAB, phbB* and *OXCT* in butanoate metabolism. Further, we validated these genes using qRT-PCR on our newly collected samples (CHN2 cohort, Data S7). Consistent with metagenomic analysis, genes of butanoate metabolism were enriched in adenoma samples, such as *bdhB* and *gctB* (Fig. 6c, Fig. S7). Conversely, genes of quorum sensing (*fusK*) and purine metabolism (*ureC, purL*) were decreased in adenomas (Fig. 6c, Fig. S7). Notably, three of the identified SNV biomarkers, SNV 1069713 of *R. bicirculans*, SNV 2604473 of *F. prausnitzii* (61481), and SNV 222146 of *B. vulgatus* were located in genes of purine metabolism, such as SNV 1069713 in the differential gene of *purL* (Data S6). Taken together, these analyses revealed the functional connections among the multimodal microbial signatures.

## Discussion

This integrated analysis provided a comprehensive multimodal view of adenoma-associated microbial signatures, including microbial multi-kingdom compositions, functional profiles and microbial SNVs. We systematically assessed their performances as non-invasive biomarkers for CRC early detection at pre-cancerous adenoma stage. Diagnostic models constructed with multi-kingdom species and genes achieved AUC values of 0.75 and 0.74, respectively. Particularly, fugal species showed superior distinguishing capabilities compared with species from other kingdoms. Meanwhile, the SNV-based diagnostic model displayed the highest accuracy (AUC=0.89) in distinguishing adenoma from control, the sensitivity and specificity of which were validated with three external adenoma cohorts. In addition, altered gene abundances in quorum sensing, purine and butanoate metabolism were observed in adenoma patients, and were further validated via qRT-PCR.

Previous efforts for microbial early detection of CRC focus on bacterial species. With 16S rRNA gene sequencing data, adenoma-specific bacterial biomarkers achieved an AUC of 0.80^12^. However, with WMS data in the current study, the diagnostic model based on bacterial species achieved a relatively low AUC of 0.66 (Fig. 4b). Similarly, a WMS study by Thomas et al.^9^ reported a low AUC with bacterial species. The discrepancy on the diagnosis power of bacteria species between 16S and WMS approaches could be attributed to insufficient coverage of the microbiome by the WMS approach. Recently, there has been an increasing interest in the roles of fungal kingdom in CRC^15,26^. We and other researchers have shown that the gut mycobiome displayed promising potentials in the early detection of CRC^18,19^. Taking advantage of the multi-kingdom-, and functional-resolution information in WMS data, the current study was able to compare the efficacies of multimodal signatures for the early screening of CRC. Notably, the diagnostic accuracy of models constructed with fungal species (AUC=0.71, Fig. 4b) was higher than those with bacteria and other kingdoms. And the combination of multi-kingdom species displayed superior diagnostic capability with an AUC of 0.75 (Fig. 4b). Therefore, these results highlight the importance of non-bacterial species in CRC early screening, especially the previously overlooked fungal kingdom. In addition, reproducible performance of gene-level architectures across cohorts have been reported in disease screening^20^. With the diagnostic model based on KO genes, we achieved an AUC of 0.74 for diagnosing adenoma, similar to that achieved with multi-kingdom species models. Interestingly, the combination of KO genes and microbial species improved the diagnosis power with an AUC of 0.78 (Fig. 4f), indicating complementary values for species and genes in detecting adenomas.

One remarkable finding in our study is that diagnostic models constructed with microbial SNV biomarkers achieved the highest AUC. First, classification models constructed with SNVs from single strains outperformed both multi-kingdom species-and gene-models. Further, the classification model constructed with SNVs from multiple strains achieved the highest accuracy (AUC=0.89, Fig. 4e). And then, a minimal set of 36 SNVs from 12 strains achieved an AUC of 0.87 with high robustness and specificity for adenoma, which could serve as a cost-effective non-invasive biomarker panel for CRC early screening. Although the bacterial kingdom did not display strong diagnostic ability in the species-level biomarkers, the abundant bacterial species enabled a more in-depth perspective towards the gut microbiome as in the high-resolution SNVs^27^. The observation that most SNV biomarkers belong to non-differential species, such as *F. prausnitzii* and *E. coli* (Fig. 5a, Data S6) further highlights the importance of analysing the genetic variations and their prominent roles as novel biomarkers for early detection at pre-cancerous stage of CRC.

Gut microbiome influence host homeostasis in multiple ways^28^, and the disruption of this harmonious interaction affects the initiation and progression of CRC^1^. Quorum sensing, a way of cell-to-cell communication, plays an important role maintaining the healthy gut microbial state through small molecules, such as autoinducers^29^, enabling microbial populations to efficiently synchronize microbial density and behavior with the surrounding environment like multicellular organisms. Here, damaged quorum sensing function in the microbiome of adenoma was observed, consistent with extensively altered intra-kingdom and fungal-bacterial interactions in patients with adenoma, suggesting impaired gut homeostasis (Fig. 2c, Fig. 6a). This hypothesis was further supported by the associations detected between quorum sensing pathway and bacterial- and fungal-biomarker species (Fig. 6b), and the decreased representations of two autoinducer receptors, *luxU* and *fusK*, in the microbiome of adenoma from both metagenomic analysis and qRT-PCR (Fig. 2c, 6c). Meanwhile, the purine metabolism was impaired in the microbiome of adenoma patients, with two of the key genes, *purL* and *ureC* exhibiting decreased abundances in adenoma (Fig 6c). Notably, three SNV biomarkers were located in the differential purine metabolism pathway, one of which was in *purL*, suggesting potential microbiome-driven mechanisms of impaired purine metabolism in the pathogenesis of adenoma. Abnormal purine level has been associated with microbial dysfunctions and cancer progression^30,31^. Here our results highlighted several ways that the microbiota of adenoma may impact the purine metabolism. On the other hand, adenoma patients were observed with increased butanoate metabolism capability. This is consistent with our previous observation of increased abundance of butanoate metabolizing genes in CRC patients compared to controls^19^, and supports an essential role of butanoate metabolism in the development of CRC^32^. This was further supported by the increased abundances of key genes in butanoate metabolism, *bdhAB* and *gctB*, via qRT-PCR. *F. prausnitzii*, one of the most abundant species in human intestines, is a major producer of butanoate^25^, and also possesses the most contributing features of the best-performing SNV model (Data S6). Taken together, these specific multimodal alterations related to microbiota metabolism and quorum sensing and their close relationships may illuminate future efforts to unravel the convoluted pathological mechanisms of adenoma and CRC, and provide reasonable explanations for the outstanding performances of the microbial biomarkers in detecting adenoma.

Collectively, we uncover comprehensive adenoma-associated microbial alterations and reveal the outstanding potential of microbial SNVs as a novel non-invasive tool for CRC early screening. In addition, we propose potential pathological mechanisms for adenoma based on the extensive alterations in the multimodal microbial interactions and the associations among the microbial biomarkers of different modalities, and of different kingdoms.

## Materials and Methods

### Patient recruitment and sample collection

Two in-house cohorts collected in Shanghai, China, were included in this study. Firstly, we collected WMS data from CHN1 cohort, including 40 adenoma patients and 47 healthy controls^45^. Besides, fecal samples were collected from CHN2 cohort, which was newly recruited from the Shanghai Tenth People’s Hospital of Tongji University, containing 29 adenoma patients and 30 healthy controls. Written informed consent was obtained from each subject before data and biospecimen collection. Patients were recruited at initial diagnosis with no reception of any treatment before fecal sample collection. Patients with hereditary colorectal syndrome, or with a previous history of colorectal disorder, were excluded. This study was approved by the Ethics Committee of the Shanghai Tenth People’s Hospital of Tongji University (ethical approval No. 20KT863).

### Public data collection

To conduct an integrative meta-analysis, we further collected published fecal WMS data of six cohorts consisting adenoma patients and healthy controls covering samples from five different countries. The metadata were manually curated from relevant original publications. Only colorectal adenoma samples and healthy controls were included for downstream meta-analysis, while samples of CRC patients were used to validate the specificity of microbial biomarkers, along with samples from CD and UC patients.

### Study design

In total, WMS data of 750 samples of adenoma patients and healthy controls from seven geographically distinct cohorts were included in the meta-analysis of this study. To identify and validate global microbial biomarkers across cohorts, four cohorts were set as the discovery dataset, including cohorts AUT, FRA, ITA, and JPN, while cohorts USA1, USA2, and one in-house cohort from China (CHN1) were used as the validation dataset. Meanwhile, samples of CRC, CD and UC patients were used to externally estimate the specifity of microbial biomarkers and classification models against adenoma. Further, newly collected samples from CHN2 cohort were used to perform qRT-PCR validations of key genes.

### Sequencing data preprocessing

KneadData (http://huttenhower.sph.harvard.edu/kneaddata, V0.6.0) was used to perform quality control on sequencing data. First, low-quality reads were removed using Trimmomatic (SLIDINGWINDOW:4:20 MINLEN:50 LEADING:3 TRAILING:3). Remaining reads were then mapped to the mammalian genomes (hg38, felCat8, canFam3, mm10, rn6, susScr3, galGal4 and bosTau8; UCSC Genome Browser), 21288 bacterial plasmids (NCBI RefSeq database accessed on January 2020), 3890 complete plastomes (NCBI RefSeq database accessed on January 2020) and 6093 UNiVec sequences (NCBI RefSeq database accessed on January 2020) by bowtie2 (V.2.3.5) to remove sequences of human and laboratory contaminations^33^.

### Microbial multi-kingdom assignment

A customized reference database comprising 18756 bacterial, 359 archaeal, 9346 viral reference genomes from the NCBI Refseq database (accessed on January 2020), and 1094 fungal reference genomes from the NCBI Refseq database, FungiDB (http://fungidb.org) and Ensemble (http://fungi.ensembl.org, accessed on January 2020) was built for taxonomic assignment of the sequencing reads. Taxa were assigned to sequencing reads using Kraken2, an improved taxonomic classification system using exact K-mer matches^34^. Further, Bracken was used for taxa abundance estimation based on Kraken2 results^35^. Read counts of species were converted to relative abundances and only those with relative abundances more than 0.1% in at least 10% of samples and presented in at least three cohorts were subjected to further analysis.

### Functional annotation

For microbial functional profiling, high-quality reads were assembled into contigs using Megahit (V1.2.9) and only contigs longer than 500-bp were selected for downstream analysis. Microbial genes were predicted by Prodigal (V2.6.3) via the metagenome mode (-p meta). Then a non-redundant microbial gene reference was constructed with CD-HIT using a sequence identity cut-off of 0.95, and a minimum coverage cut-off of 0.9 for the shorter sequences. The reference was annotated using EggNOG mapper (V2.0.1) based on EggNOG orthology. CoverM (https://github.com/wwood/CoverM, V4.0) was used to estimate gene abundances by mapping high quality reads to reference sequences and to calculate the coverage of genes in the original contigs. The relative abundances of KEGG orthologous (KO) groups or pathways were calculated by summing the relative abundances of genes annotated to the same KOs or pathways.

### SNV calling

Metagenomic Intra-Species Diversity Analysis System (MIDAS, V1.3.2) was used to perform microbial SNV annotation^36^. Briefly, a customized reference genome database including 28 strains with sufficient coverage, which is defined as the percentage of genomes from cellular organisms in a community that have a sequenced representative in the reference database (>3X as default parameter) in at least 10% of all samples, was constructed. Then, the WMS reads were mapped to the database for SNV calling. Subsequently, the SNV profiles of each sample were merged, and only bi-allelic positions were chosen. Other parameters were in accordance with the preset option “— core_snps”.

### Integrated analysis of microbiome-derived differential signatures

xMarkerFinder, an integrated workflow designed to address the cross-cohort heterogeneity of human microbiome, was employed in this study^37^. This workflow mainly comprises differential signature identification, model construction, model validation, and biomarker interpretation, as detailed in the following paragraphs.

### Cross-cohort differential signature identification

Firstly, MMUPHin (V1.4.2) was used for the identification of signatures that are differential across cohorts with respect to combined phenotypes^38,39^. Respective regression analyses in individual cohorts were performed and then aggregated with established fixed effect models to test for consistently differential signatures between adenoma and control samples with “cohort” set as the main batch and demographic indices, including gender, age and BMI, as covariates. Signatures with p values less than 0.05 were identified as differential signatures and used as input for downstream feature selection procedure.

### Feature selection

Based on multimodal differential signatures, Triple-E, a three-step feature selection procedure implemented in xMarkerFinder, was employed to identify candidate biomarkers. The first step was feature effectiveness evaluation, aiming to select individual features with discriminative power (AUC > 0.5) as effective features. Next, collinear feature exclusion was performed to exclude highly-correlated features and features with absolute values of correlation coefficient less than 0.7 were considered as uncorrelated-effective features and selected for the next step, recursive feature elimination for the identification of the panel of optimal biomarkers with the highest predictive capability. This is achieved by repeat modeling starting with all features and recursively removing the weakest feature for model construction per loop to obtain the best panel with the highest cross-validation AUC value.

### Model construction and optimization

The optimal biomarkers were then used to construct RF models with stratified five-fold cross-validation to avoid overfitting. Further, to optimize the diagnostic RF models, hyperparameters, including the number of estimator trees, the maximum depth of the trees, the numbers of features per tree, and the maximum samples were tuned using bayesian-optimization (V1.2.0) package. Finally, with the selected optimal biomarkers and the best combination of hyperparameters, the best-performing RF models were constructed.

### Evaluation of the biomarkers’ robustness and generalization

To test the robustness and generalization of identified optimal biomarkers among geographically distinct cohorts, we performed cohort-to-cohort and LOCO validation as described in our previous study^12^. For cohort-to-cohort validation, diagnostic models were trained on the profile of one single cohort and validated on the profile of each of the remaining cohorts, respectively. For LOCO validation, one single cohort was set as the validation dataset while all other cohorts were pooled together as the discovery dataset. Further, four independent cohorts were used to externally validate the robustness of identified optimal biomarkers, as well as the optimized SNV diagnostic model.

### The specificity of adenoma predictive biomarkers

To avoid false positives in clinical diagnosis, we evaluated the disease specificity of the best panel of microbial biomarkers for adenoma by examination of their performances in discriminating non-adenoma diseases from controls. These non-adenoma diseases included CRC (386 cases and 439 controls), CD (68 cases and 34 controls), and UC (53 cases and 34 controls).

### Co-abundance analysis of differential multi-kingdom species

To investigate the associations among differential multi-kingdom species, we employed SparCC^40^, a widely used approach for estimating correlations with compositional data, to construct the microbial network in different disease status. Associations among differential multi-kingdom species were inferred with 50 iterations, after which the statistical significance was calculated with 1000 permutations. Correlations with |r| > 0.5 and p-value <0.05 were regarded as moderate correlations and were included in downstream analysis. Network was visualized with Gephi (V0.9.2).

### Associations between microbial species and function

To further explore the potential associations between multimodal signatures, HAllA (V 0.8.18), a computational method to find multi-resolution associations in high-dimensional heterogeneous datasets, was applied to evaluate the associations between differential pathways and multi-kingdom species^41^.

### qRT-PCR validation

To quantify the abundances of key genes, qRT-PCR analysis was performed in triplicates on newly collected samples of CHN2 cohort (29 adenomas and 30 controls). The genomic DNA was extracted with the TIANamp Stool DNA Kit (Cat# 4992205, TIANGEN) according to the manufacturer’s instructions. We used the primers in Data S8 for candidate genes, and standard primers F515 and R806 for 16S rRNA. To perform the qRT-PCR reaction, the final primer concentration was diluted to 0.2 μM including 10 ng of genomic DNA in a 10 μl final reaction volume with the SYBR Green qPCR Mix (Thermo Fisher Scientific). The adopted qRT-PCR program was as follows: predenaturation at 95 °C for 10 min; denaturation at 95 °C for 15 s and annealing at 60 °C for 60 s for 40 cycles; followed by a melting curve analysis. The qRT-PCR result was quantitated by calculate 2^-ΔΔCt^ values between candidate genes and 16S Ct values as the relative expression level.

### Statistical analysis

Considering the sparsity of microbial data, nonparametric Wilcoxon tests and a threshold of 0.05 in *P* values were used unless stated otherwise. PERMANOVA test was performed to quantify the contributions of the subjects’ physical variables to multimodal microbial profiles using R (V4.0.5) “vegan” (V2.5.7) package with 999 permutations^42^. Alpha diversity metrics, including Shannon and Simpson Index, and beta diversity based on Bray-Curtis distance of taxonomic and functional profiles were calculated. Differences between groups were then estimated with MaAsLin 2 (V1.4.0) and PERMANOVA, respectively^43^. All statistical and bioinformatics analyses were implemented using R (V 4.0.5) and Python (V 3.8.5).

## Data and code availability

All processed data for this work are available upon request. xMarkerFinder is provided at https://github.com/tjcadd2020/xMarkerFinder.

## Abbreviations

AUC: area under the ROC curve;
BMI: body mass index;
CD: crohn’s disease;
CDS: coding sequence;
CRC: colorectal cancer;
CTC: circulating tumor cells;
ctDNA: circulating tumor DNA;
IBD: inflammatory bowel disease;
IGR: intergenic region;
KO: KEGG orthology;
MIDAS: Metagenomic Intra-Species Diversity Analysis System;
MMUPhin: Meta-analysis Methods with a Uniform Pipeline for Heterogeneity in Microbiome Studies;
NAFLD: non-alcoholic fatty liver disease;
PCoA: principal coordinate analysis;
PERMANOVA: permutational multivariate analysis of variance;
qRT-PCR: quantitative real-time PCR;
RF: random forest;
SNV: single-nucleotide variants;
UC: ulcerative colitis;
WMS: whole metagenome sequencing.

## Acknowledgments

The authors would like to thank all the researchers for generously sharing their sequencing data included in this study. We acknowledge funding from the National Natural Science Foundation of China (82170542 to RZ, 92251307 to RZ, 91942312 to ZL, 81630017 to ZL, 32200529 to DW, 82000536 to NJ), the National Key Research and Development Program of China (2021YFF0703700/2021YFF0703702 to RZ), and Guangdong Province “Pearl River Talent Plan” Innovation and Entrepreneurship Team Project (2019ZT08Y464 to LZ). The funders had no role in study design, data collection and analysis, decision to publish, or preparation of the manuscript.

## Author contributions

NJ, RZ, ZL and LZ conceived and designed the study. WG, RS, SG, ZF, DW, XG, and NJ performed the data analysis and model construction. RS, ZF, and XG collected the fecal sample and performed the experimental validation. WG and NJ wrote the manuscript. LZ, ZL, RZ and NJ reviewed and edited the manuscript. All authors read and approved the final manuscript.

## Supplementary files

**Supplemenraty Figures S1–S7.**

**Figure. S1.**
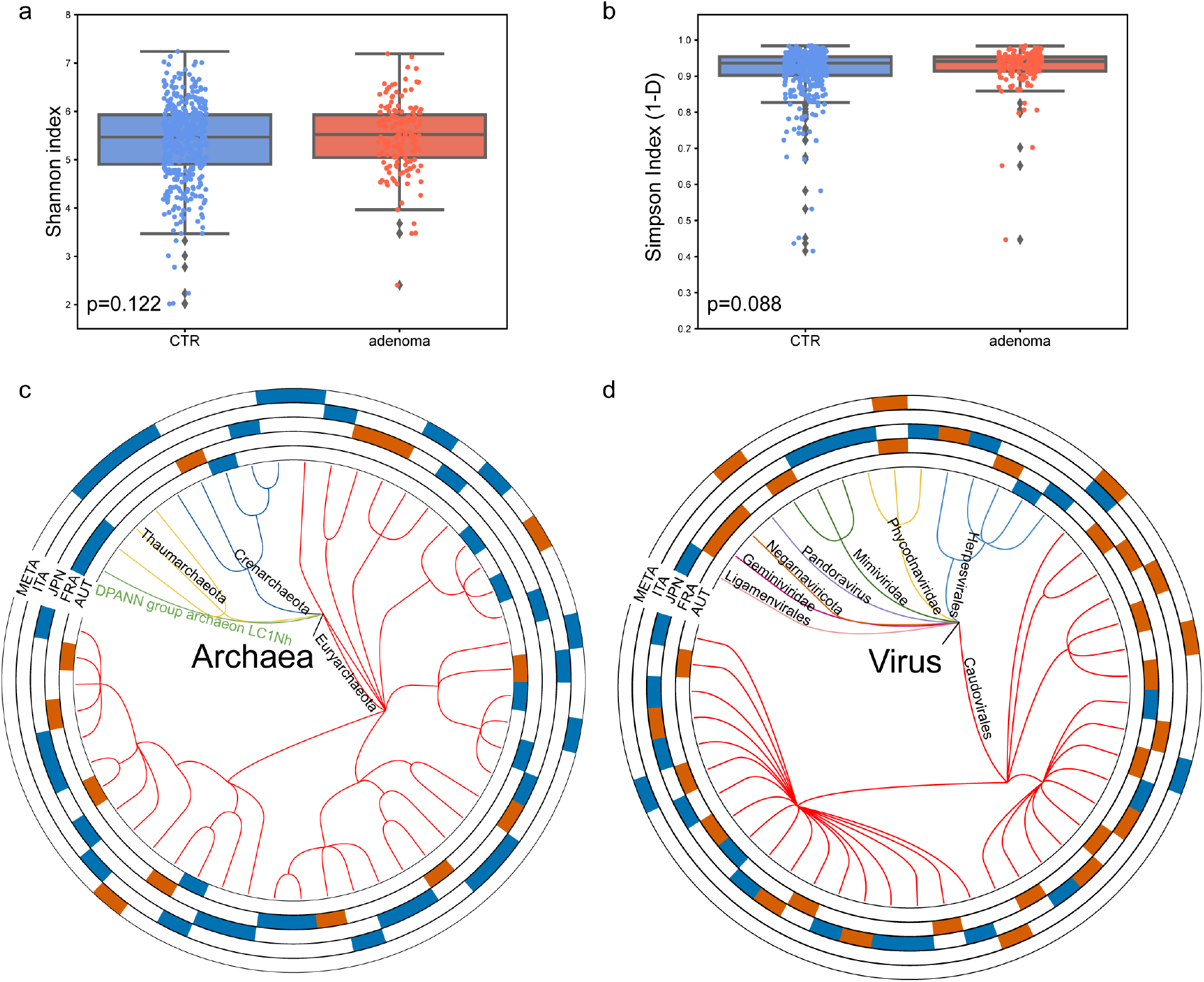
Adenoma-associated microbial taxonomic alterations. **a,b**, Alpha diversity as measured with the (**a**) Shannon Index and (**b**) Simpson Index of multi-kingdom taxonomic profiles was computed in all samples. P-values between two groups were calculated via MaAsLin2. **c,d**, Phylogenetic trees showing the union of different archaeal (**c**, 50 in total) and viral species (**d**, 50 in total). The outer circles are marked for significant differential species (*P* < 0·05) in each cohort identified via Maaslin2 and in the meta-analysis identified via MMUPHin (META ring) with orange for increased species and blue for decreased species.

**Figure. S2.**
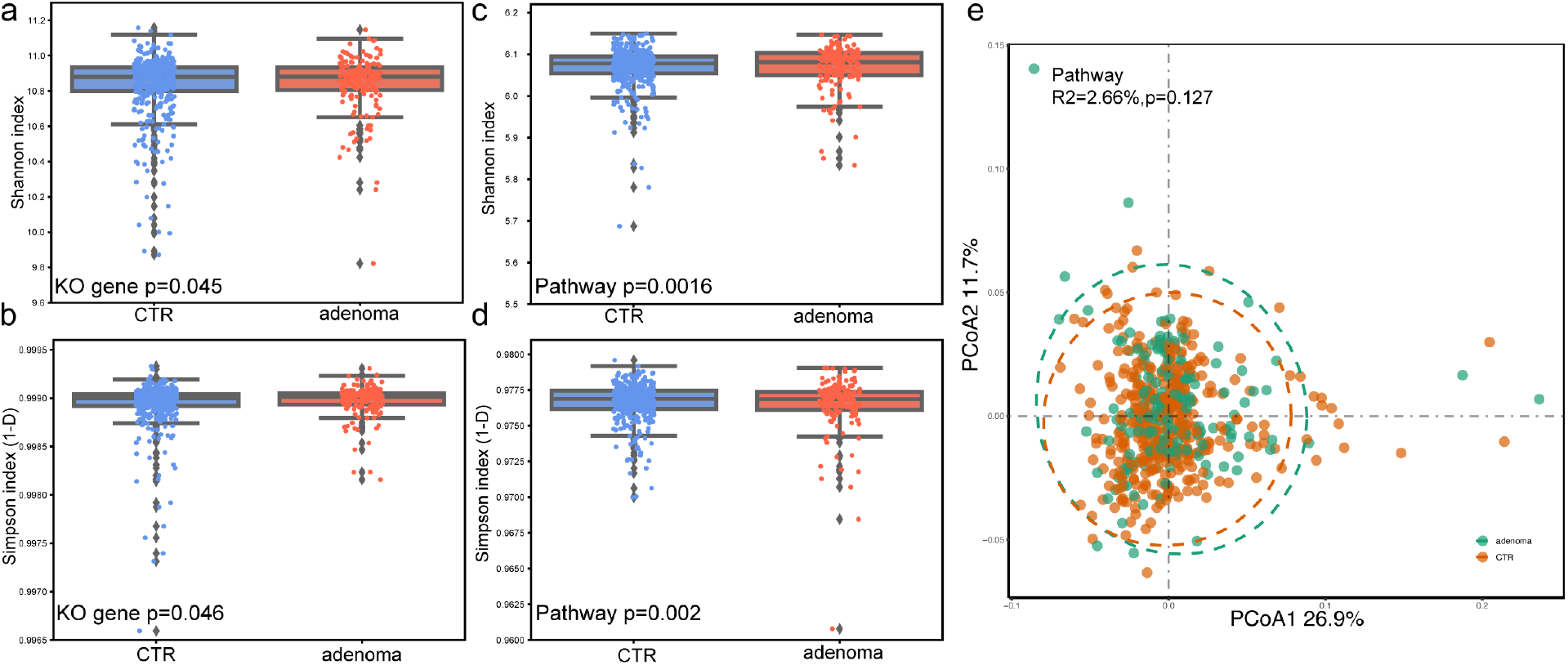
Alpha and beta diversity in microbial functional profiles between adenoma patients and controls. **a,b**, Alpha diversity as measured with the (**a**) Shannon Index and (**b**) Simpson Index of KO genes was computed in all samples. *P* values between two groups were calculated via MaAsLin2. **c,d**, Alpha diversity as measured with the (**c**) Shannon Index and (**d**) Simpson Index of functional pathways was computed in all samples. *P* values between two groups were calculated via MaAsLin2. **e**, Principal coordinate analyses (PCoA) of functional pathways showing no significant differences between adenoma patients and healthy controls. *P* values of beta diversity based on Bray-Curtis distance were calculated with PERMANOVA test. Each point in the PCoA plots represents a sample and the colors of points represent different groups.

**Figure. S3.**
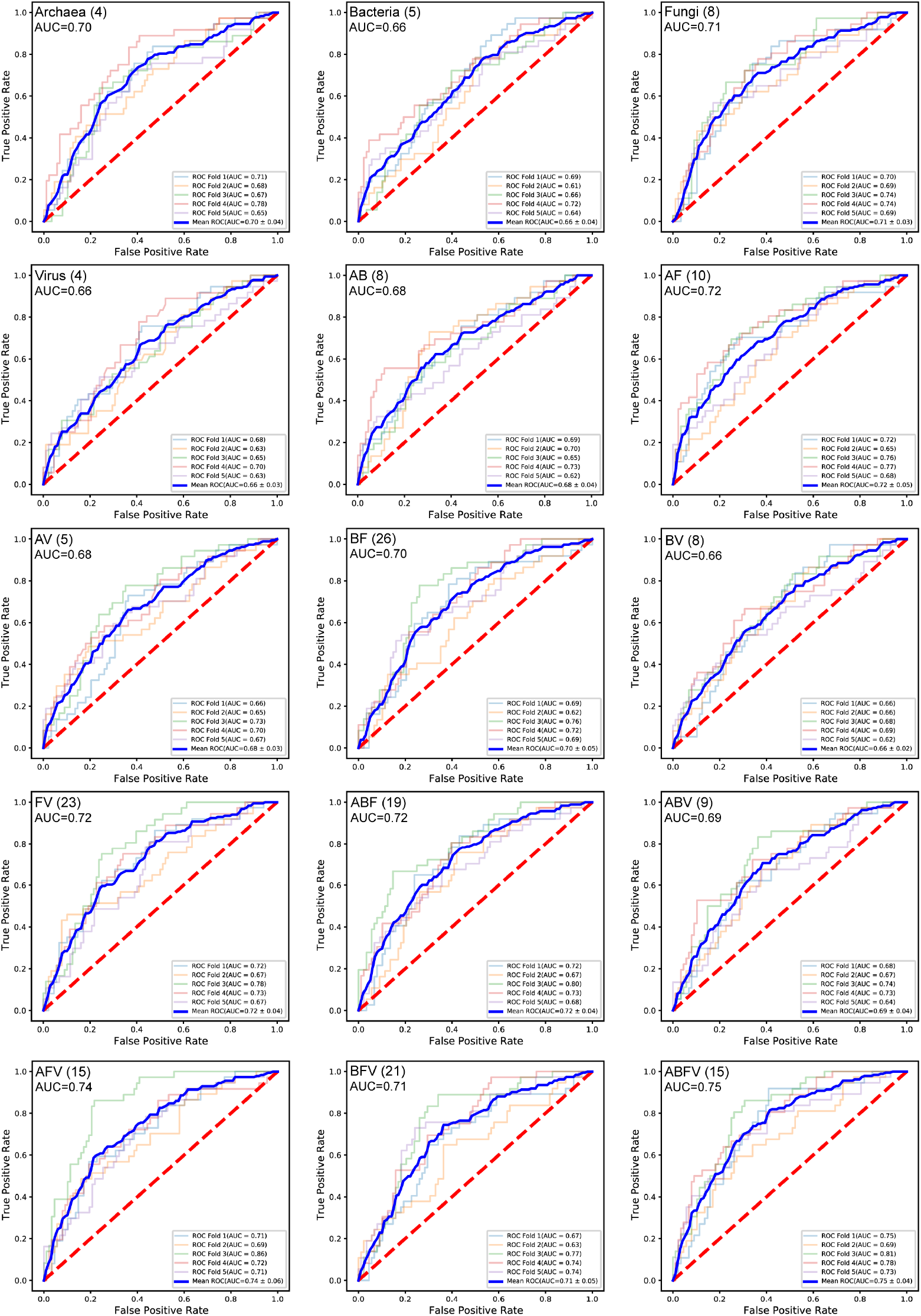
Performances of models based on multi-kingdom features. Receiver operating characteristic (ROC) curves of the optimized models constructed with multi-kingdom features. The numbers of used features and mean AUC were shown.

**Figure. S4.**
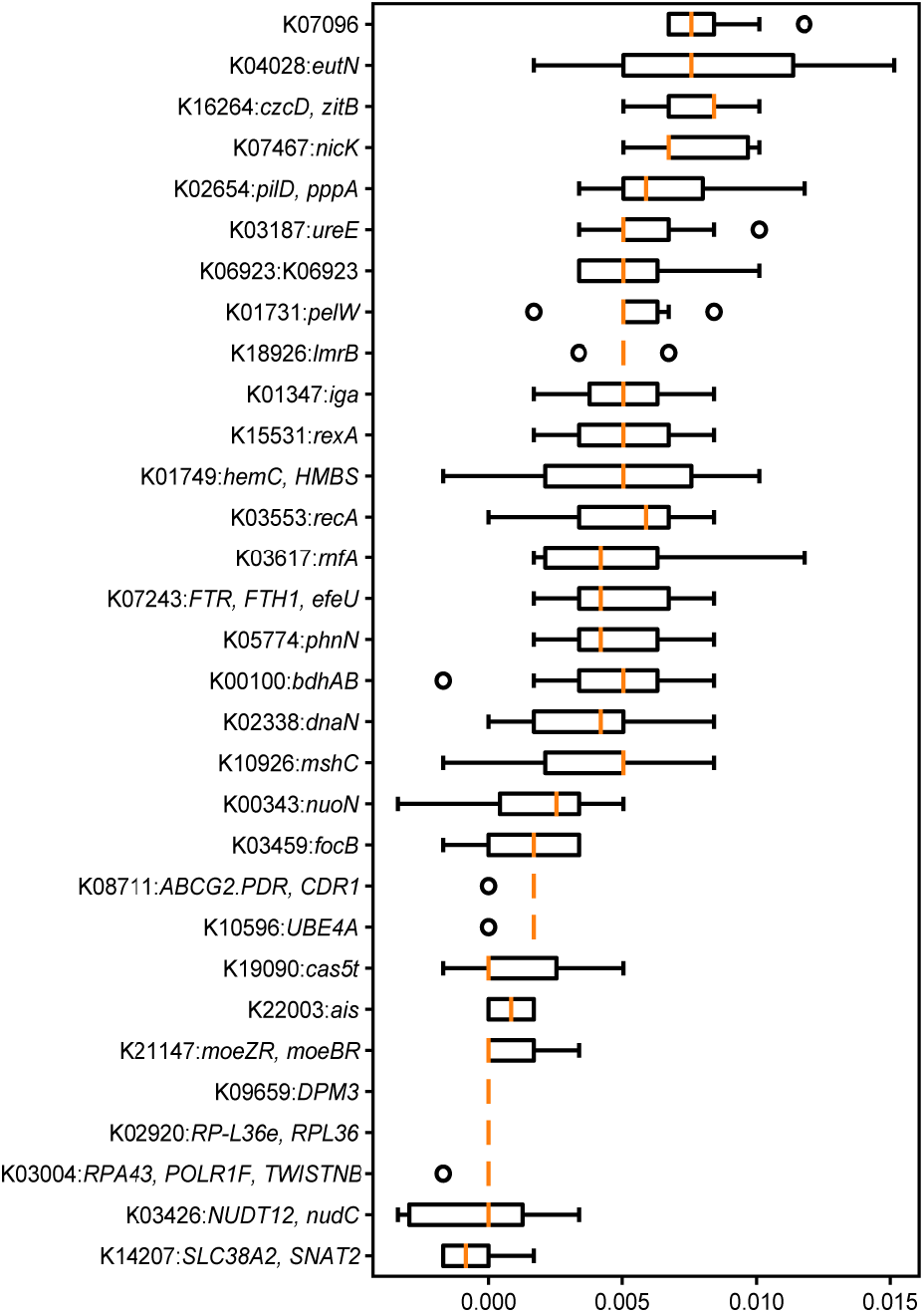
Feature importances of the optimal KO gene model. Permutation feature importances of the optimized model constructed with KO genes.

**Figure. S5.**
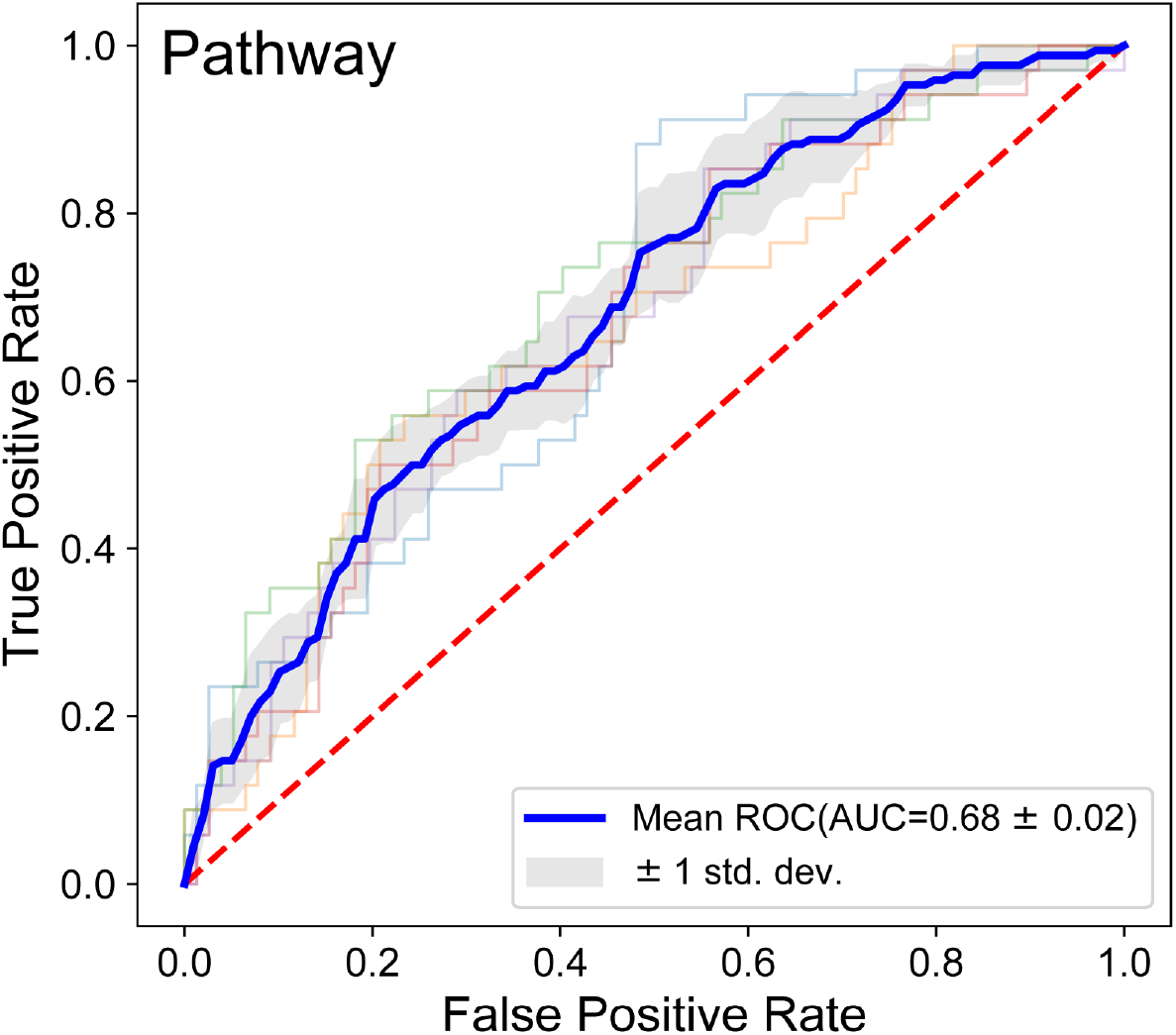
Performance of the diagnostic model based on functional pathways. Receiver operating characteristic (ROC) curve of the optimized models constructed with pathway-level biomarkers. Mean AUC and standard deviation of stratified 5-fold cross-validation were shown.

**Figure. S6.**
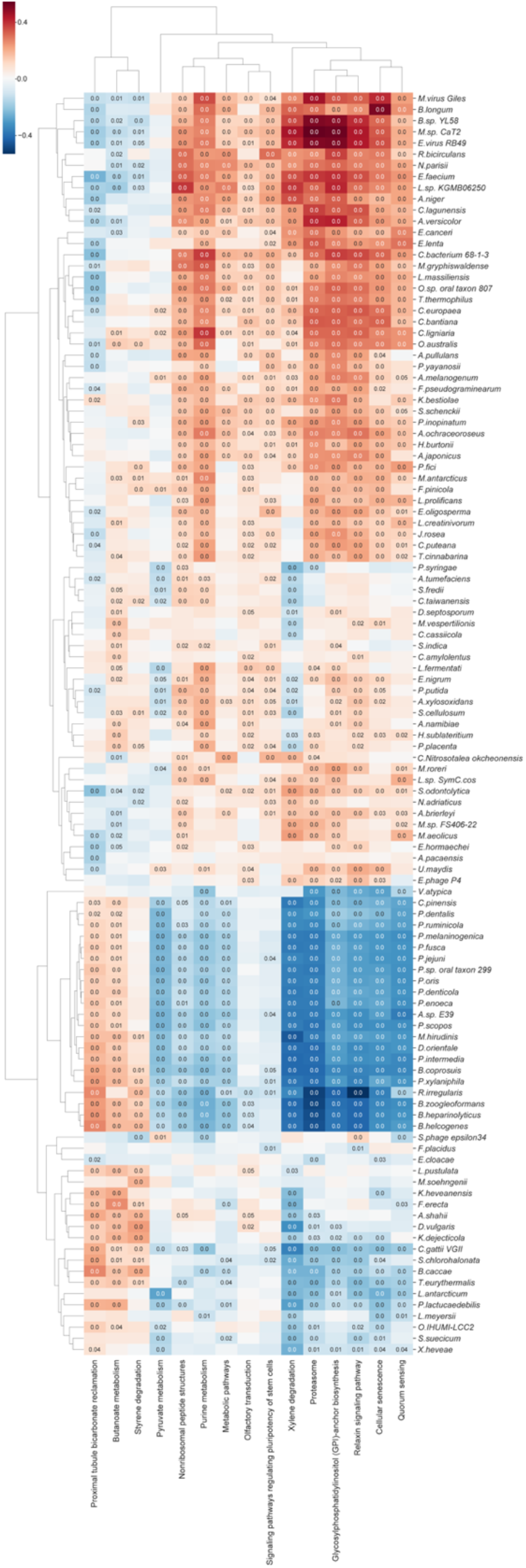
Associations between differential pathways and differential species. Hierarchical All-against-All association testing (HAllA) between differential pathways and differential species. Significant associations between pathways and species were plotted after BH-FDR correction. Red and blue cells indicate positive and negative correlations, respectively, and color intensity of the heatmap and the numbers in the cells indicate significant pairs of associations.

**Figure. S7.**
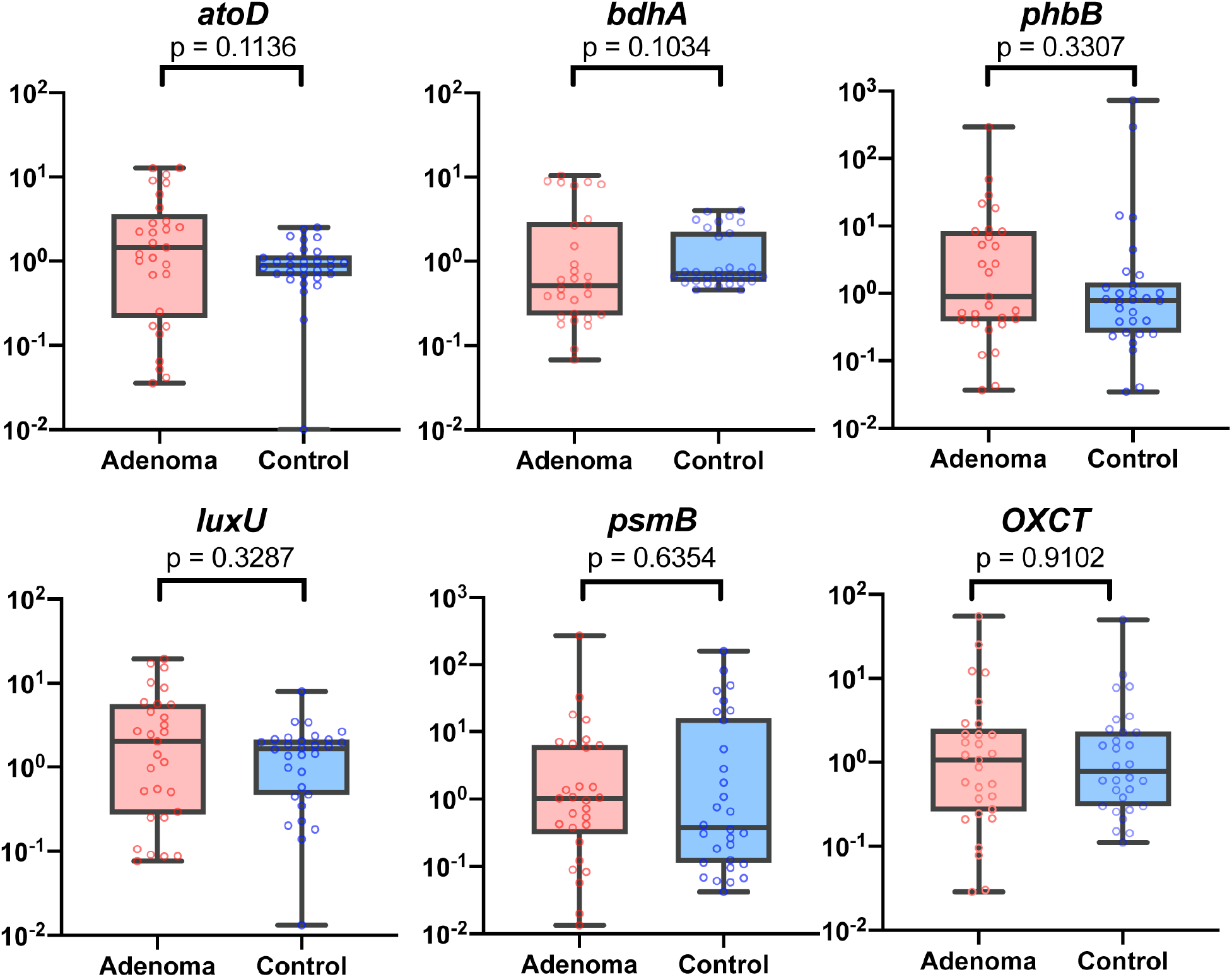
qRT-PCR results of key genes. qPCR results showing the expression of key genes. *P* values were computed using a two-sided Wilcoxon rank-sum test.

**Supplementary_Data.xlsx contains Data S1-S8.**

**Data S1.**Characteristics of the patients and controls included in metagenomic analyses

**Data S2.**Differential multi-kingdom taxonomic signatures across cohorts between adenomas and

controls

**Data S3.**Differential gene signatures across cohorts between adenomas and controls

**Data S4.**Differential pathway signatures across cohorts between adenomas and controls

**Data S5.**SNV profiles of commonly detected strains in the discovery dataset

**Data S6.**Site-specific profiles of biomarkers in the best-performing SNV model

**Data S7**. Characteristics of the patients and controls enrolled for qRT-PCR validations

**Data S8**. Primers for qRT-PCR validations

